# A mechanically actuated lung microvascular model reveals that breathing-like deformation enhances tumor cell extravasation

**DOI:** 10.64898/2026.09.17.752385

**Authors:** Marie A. Floryan, Roland M. Stoll, Ingmar Fleps, Rachael N. Coates, Roger D. Kamm, Elena Cambria

**Affiliations:** Department of Biological Engineering, Massachusetts Institute of Technology, Cambridge, MA, USA; Department of Mechanical Engineering, Massachusetts Institute of Technology, Cambridge, MA, USA; Independent researcher

**Author notes:** These authors contributed equally (Marie A. Floryan, Roland M. Stoll).

## Abstract

The lung is a common site of metastasis and is continuously deformed by breathing. Whether physiological breathing mechanics influence tumor cell extravasation is unknown. To address this, we developed a mechanically actuated microphysiological system combining perfusable lung-specific microvascular networks with breathing-like cyclic deformation. Under static conditions, lung-specific networks exhibited more lung-like vascular features and higher tumor cell extravasation rates than generic networks. The mechanically actuated device reproduced physiological tissue and vascular deformation, while 24 h of cyclic actuation did not alter vessel viability, permeability, or morphology compared with unactuated controls. Pre-actuation for 24 h, followed by tumor cell perfusion under static conditions, did not alter subsequent extravasation, whereas actuation initiated immediately after perfusion significantly increased extravasation compared with unactuated controls. This biomimetic platform enables direct investigation of how breathing-like vascular deformation influences metastasis and provides a versatile tool for studying vascular mechanobiology in the lung and other mechanically active organs.

## INTRODUCTION

The lung is a highly dynamic organ that undergoes continuous cyclic deformation during breathing. The alveoli are surrounded by a dense capillary network embedded within extracellular matrix containing interstitial lung fibroblasts, all of which are mechanically deformed during lung expansion and recoil. Physiological breathing has been associated with linear alveolar strains of approximately 4-10% [1–6] and occurs at a frequency of approximately 0.2 Hz (12 breaths/min) at rest [7]. These breathing-induced mechanical strains are sensed by lung cells and contribute to lung and vascular homeostasis, while changes in strain magnitude can alter lung endothelial function [8–10]. The lung is also a common site of metastasis, including for breast cancer [11, 12]. Circulating tumor cells reaching the lung can become physically trapped within the small lung capillaries before extravasating into the surrounding tissue [13]. The highly interconnected architecture of the alveolar capillary network allows blood flow to be redistributed around local obstructions, limiting pressure buildup behind an occluded capillary segment and potentially favoring the retention of lodged tumor cells [14]. Although mechanical ventilation has been shown to increase tumor cell invasiveness and metastatic progression [15, 16], these studies used externally imposed ventilation that can induce non-physiological lung deformation and tissue injury. Whether physiological breathing-like mechanical deformation influences tumor cell extravasation in the lung remains unknown.

Studying the role of breathing mechanics in lung metastasis is challenging with existing experimental models. In animal models, breathing cannot be stopped for prolonged periods without compromising viability, making it difficult to isolate breathing-induced mechanical deformation as an independent experimental variable. Intravital microscopy has enabled direct visualization of tumor cells within lung capillaries, but the imaged lung region is immobilized to reduce breathing-induced motion [13]. More recently, the Crystal Ribcage platform, an ex vivo mouse lung model, enabled imaging of ventilated lungs and showed that tumor cells lodged within pulmonary capillaries deform in response to changes in alveolar pressure [17]. However, tumor cell extravasation was not observed in this model. These limitations highlight the need for controlled in vitro systems that combine breathing-like mechanical deformation with perfusable lung microvasculature.

Microphysiological systems provide an opportunity to isolate and control mechanical cues. Several lung-on-chip models have been developed to reproduce breathing-like cyclic deformation, primarily using epithelial and/or endothelial cell layers cultured on deformable membranes or substrates [18–21]. Many of these systems rely on multilayer membrane-based architectures, in which a deformable membrane is sandwiched between separate microfluidic compartments. Although these systems have provided important insights into lung mechanobiology, they do not reproduce the three-dimensional, perfusable architecture of the lung microvasculature. More recently, cyclic mechanical actuation has been applied to three-dimensional vascular models, including endothelialized vessels embedded within an extracellular matrix [22, 23]. These models enable the study of vascular responses to mechanical deformation but do not reproduce an interconnected microvascular network in which circulating tumor cells can arrest and extravasate.

In parallel, self-assembled microvascular networks have been incorporated into lung tissue and disease models [24–26] and have recently been used to study tumor cell extravasation through lung-specific microvasculature [27]. These models, however, lack breathing-like mechanical actuation. A small number of recent models have begun to combine three-dimensional microvasculature with dynamic lung mechanics, including a ventilated lung model containing perfusable microvascular networks derived from human umbilical vein endothelial cells (HUVECs) [28] and a vascularized model of bronchoconstriction-induced airway remodeling [29]. The latter reproduces compressive loading associated with airway constriction rather than physiological breathing deformation. However, to our knowledge, no current model combines perfusable lung-specific self-assembled microvascular networks with physiological breathing-like mechanical actuation to investigate tumor cell extravasation.

In this study, we first generated perfusable lung-specific microvascular networks under static conditions and compared their vascular barrier function, structure, and tumor cell extravasation rate with generic microvascular networks derived from immortalized HUVECs (iHUVECs). We then designed and characterized a mechanically actuated microfluidic device with a single-layer architecture, rather than the multilayer membrane-based designs commonly used in breathing lung-on-chip systems. The device was capable of reproducing physiological breathing-like deformation. By incorporating lung-specific microvascular networks into the mechanically actuated device, we directly quantified vascular deformation and assessed the effects of cyclic actuation on vessel viability, permeability, and morphology. Finally, we used the system to determine how breathing-like mechanical actuation influences the extravasation of perfused tumor cells.

## RESULTS

### Lung-specific microvascular networks are perfusable and display distinct morphological traits

Aiming to develop a microphysiological model able to mimic human lung physiology as closely as possible, we first set out to develop microvascular networks composed exclusively of human primary cells. Human pulmonary artery endothelial cells (hPAECs) and human normal lung fibroblasts (hNLFBs) were used to form lung-specific microvascular networks in a fibrin gel within a classic three-channel microfluidic device under static conditions (**Fig. 1A**). These lung-specific networks were benchmarked against microvascular networks routinely used in the field composed of iHUVECs and hNLFBs under similar conditions (**Fig. 1A**). Both hPAEC- and iHUVEC-microvascular networks self-assembled over the course of 7 days (**Fig. 1B**), resulting in fully perfusable vessel lumina as shown by perfusion of fluorescently labeled dextran (70 kDa) (**Fig. 1C**). Dextran-perfused microvascular networks were further imaged by confocal microscopy to assess vessel permeability and morphology. While hPAEC- and iHUVEC microvascular networks exhibited similar barrier function (with an average of 3.13 x 10^-8^ cm/s for iHUVECs and 6.20 x 10^-8^ cm/s for hPAECs, **Fig. 1D**), morphology analysis revealed that hPAECs formed microvascular networks with a lower vascular projected area (**Fig. 1E**), higher vessel density (**Fig. 1F**), and shorter vessels (**Fig. 1G**) compared to iHUVEC microvascular networks. Lung-specific microvascular networks also generally exhibited vessels of smaller diameter (**Fig. 1H**) and higher branching point density (**Fig. 1I**). Overall, these results show that our lung-specific microvascular networks are perfusable and possess morphological traits reminiscent of human lung vasculature such as dense and short vessels.

**Figure 1.**
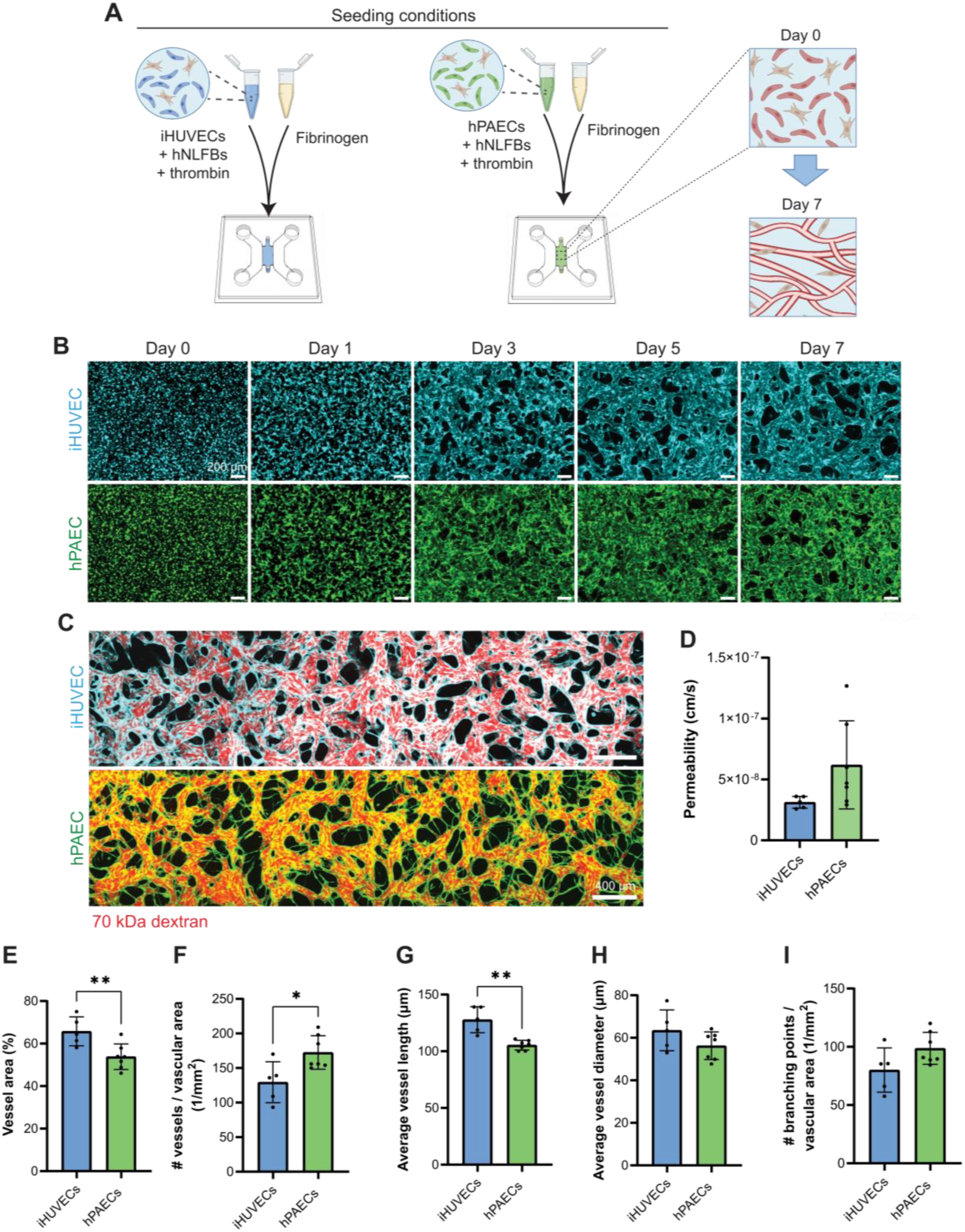
Lung-specific microvascular networks are perfusable and display distinct morphological traits. (A) Graphic describing cell culture conditions in a static microfluidic device on day 0 and an overview of the 7-day microvascular network formation process. iHUVECs = immortalized human umbilical vein endothelial cells. hNLFBs = human normal lung fibroblasts. hPAECs = human pulmonary artery endothelial cells. (B) Representative epifluorescence microscopy images of microvascular network formation with iHUVECs or hPAECs on days 0, 1, 3, 5, and 7. Scale bar is 200 μm. (C) Representative confocal microscopy images of dextran-perfused microvascular networks showing full vessel perfusability. Scale bar is 400 μm. (D) Permeability analysis of vessels formed with iHUVECs or hPAECs. n = 5-7 devices. (E-I) Morphology analysis of vessels formed with iHUVECs or hPAECs including (E) projected vessel area fraction, (F) vessel density, (G) average vessel length, (H) average vessel diameter, and (I) vessel branching point density. n = 5-7 devices. * indicates p < 0.05, ** indicates p < 0.01.

### Tumor cell extravasation rate is higher in lung-specific microvascular networks

After having characterized lung-specific microvascular networks in terms of vascular permeability and morphology, we next assessed them functionally in a tumor cell extravasation assay (**Fig 2A**). Fluorescently labeled triple-negative breast cancer cells (MDA-MB-231) were perfused into fully formed microvascular networks containing hNLFBs combined with either iHUVECs or hPAECs. 42 hours after tumor cell perfusion, microvascular networks were imaged under a confocal microscope (**Fig. 2B**) and extravasated cells were counted and normalized by the total number of tumor cells in each microfluidic device to calculate the extravasation rate. Despite having a similar total number of tumor cells per device (**Fig. 2C**), we observed a stark increase in MDA-MB-231 cell extravasation rate in hPAEC microvascular networks compared to iHUVEC microvascular networks (with an average of 8.09 % for iHUVECs and 27.46 % for hPAECs, **Fig. 2D**), suggesting the involvement of underlying lung-specific factors.

**Figure 2.**
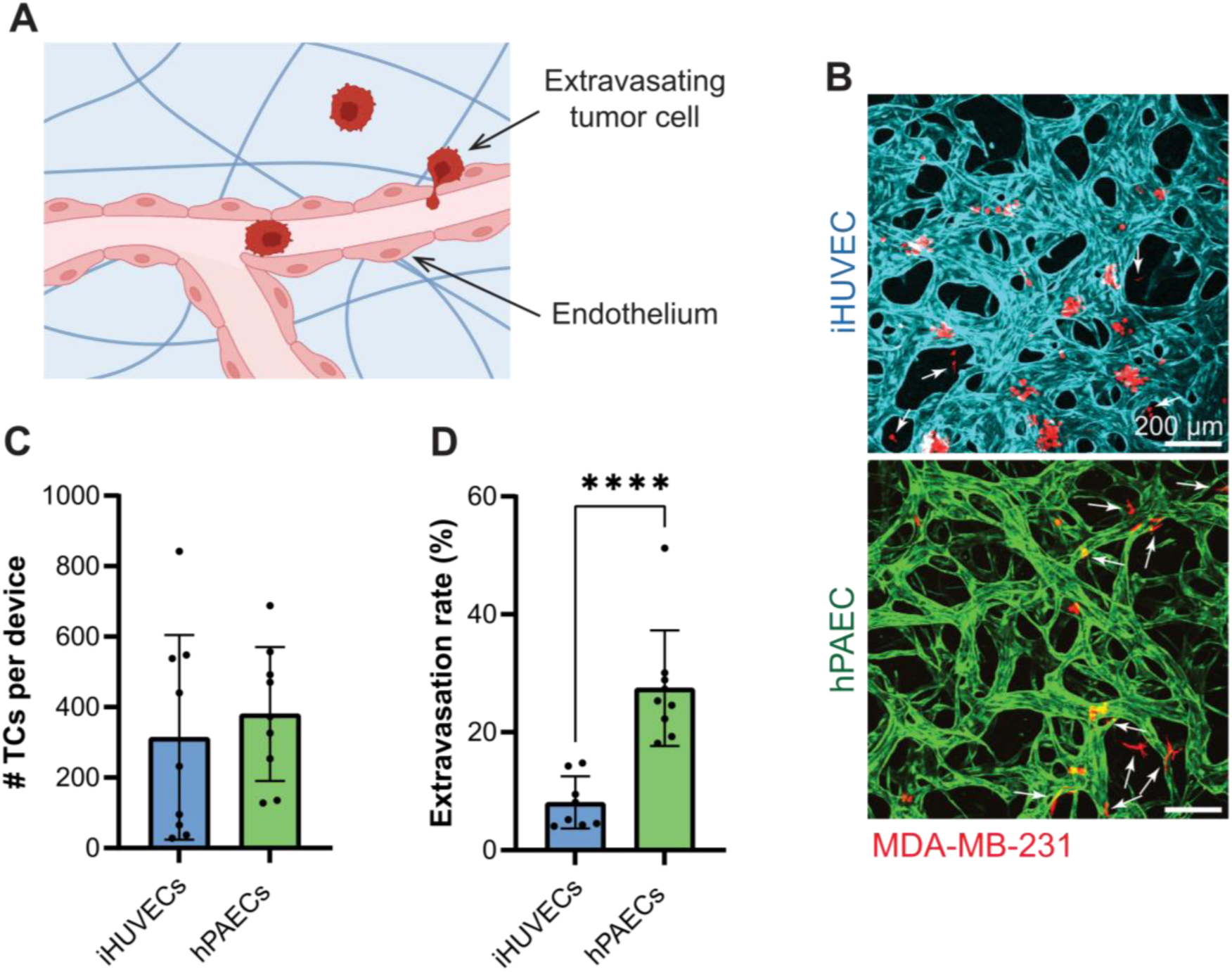
Tumor cell extravasation rate is higher in lung-specific microvascular networks. (A) Illustration describing the process of tumor cell extravasation across the endothelium. (B) Representative confocal microscopy images of microvascular networks formed with either iHUVECs or hPAECs and perfused with MDA-MB-231 tumor cells. Extravasated tumor cells are indicated by white arrows. Scale bar is 200 μm. (C-D) Tumor cell extravasation analysis showing (C) total number of tumor cells (TCs) per device and (D) tumor cell extravasation rate in microvascular networks formed with iHUVECs or hPAECs. n = 8-9 devices. **** indicates p < 0.0001.

### A mechanically actuated microfluidic device mimics physiological breathing micromechanics

In the next step, we aimed to design and fabricate a microfluidic device that could accommodate the lung-specific microvascular networks and mimic the micromechanics of the alveolus during normal breathing. At lower lung volume, the lung alveoli are less inflated and the surrounding capillaries within the interstitium display a more circular cross section (**Fig. 3A**). At higher lung volume, the alveoli expand and the capillaries are stretched along the alveolar wall and compressed radially, resulting in a reduced cross section (**Fig. 3A**) [30]. To replicate these mechanical strains, we designed a single-layer microfluidic device composed of a gel channel that can house the microvascular networks in fibrin (**Fig. 3B** and **3Ci**). In absence of traditional media channels, nutrient transport is ensured through syringes affixed to both ends of the gel channel, generating a difference in hydrostatic pressure and resulting in medium flowing along the gel channel (**Fig. 3B**). Additionally, two air channels extend perpendicularly from each side of the gel channel and are separated from the gel by two thin polydimethylsiloxane (PDMS) walls, forming a cross-shaped junction (**Fig. 3B** and **3Ci-ii**). These air channels are connected to a pneumatic actuation system that applies vacuum pressure on the thin PDMS walls and ceiling of the device, resulting in multidirectional mechanical strains in the gel that mimic the strains experienced by lung capillaries in vivo (**Fig. 3Cii**).

**Figure 3.**
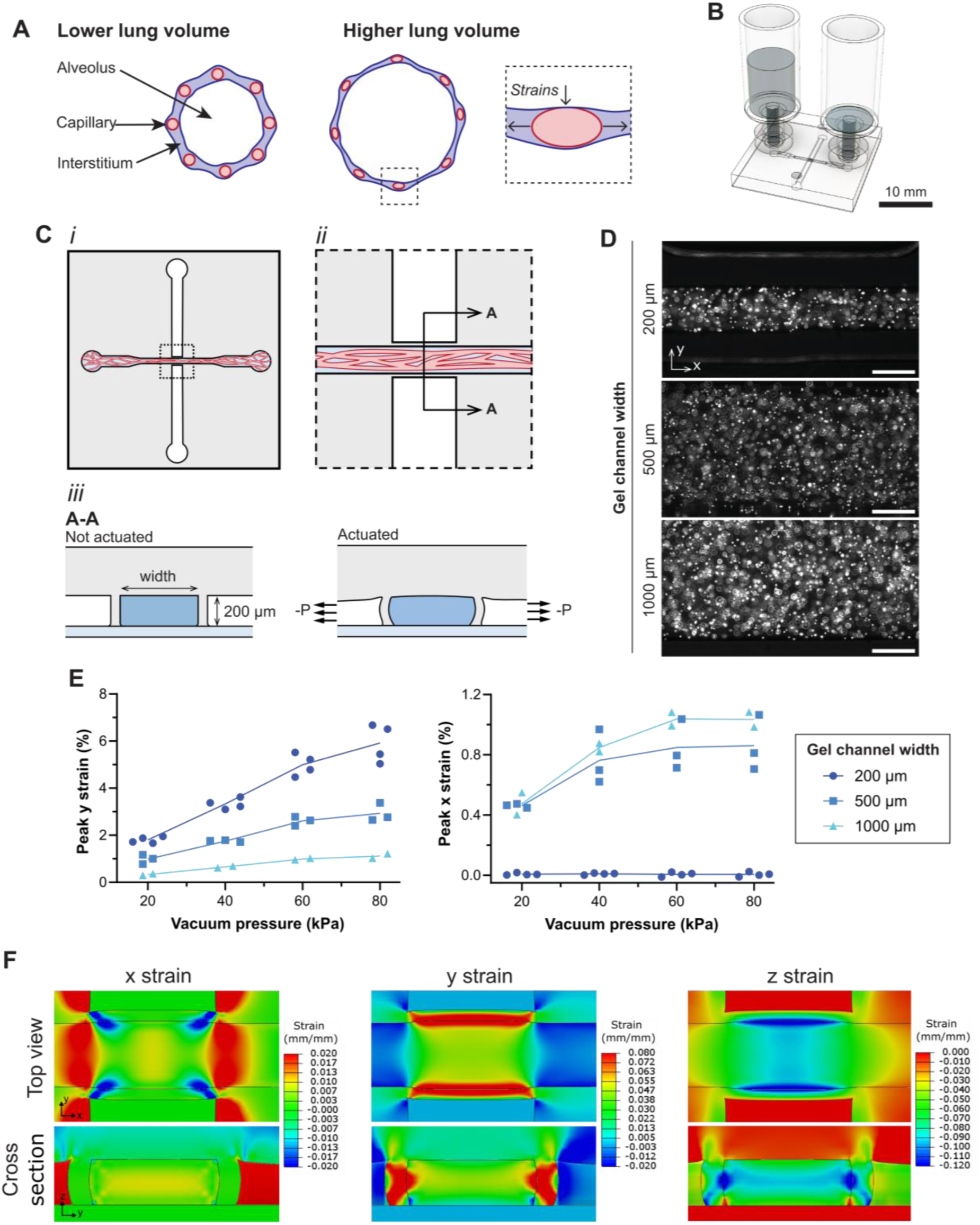
A mechanically actuated microfluidic device mimics physiological breathing micromechanics. (A) Simplified schematic illustrating alveolar and capillary deformation at lower and higher lung volumes during tidal breathing. (B) Mechanically actuated microfluidic device with two 3 mL syringes affixed to both ends of the gel channel, generating a hydrostatic pressure difference of 1 cm H_2_O and resulting in medium flow along the gel channel. Scale bar is 10 mm. (C) Top view of the mechanically actuated microfluidic device showing (i) the whole device and (ii) a magnified view of the cross-shaped junction including the gel channel with microvascular networks (red) and the two perpendicular air channels for vacuum actuation (white) separated by thin PDMS walls (gray). (iii) Cross-sectional view of the actuation region of the gel channel (blue) in the not actuated and actuated states, before and during application of vacuum pressure. (D) Representative epifluorescence microscopy images of the actuation region showing fluorescent microspheres embedded in the fibrin gel in devices with three different gel channel widths used for DIC analysis. Scale bar is 200 μm. (E) Characterization of peak y and x strains for the three gel channel widths at vacuum pressures ranging from 20 to 80 kPa, quantified by DIC. Each device was tested at all four vacuum pressures. Lines connect the mean values for each channel width across pressure conditions. n = 2-4 devices per channel width. (F) Top and cross-sectional views of the x-, y-, and z-strain distributions through the central planes of the gel, obtained by FEA for a 500-μm-wide gel channel at a vacuum pressure of 80 kPa.

To identify a design capable of generating physiological strains, we fabricated microfluidic devices with gel channel widths of 200, 500, and 1000 μm. Mechanical strains along (x strain) and across (y strain) the gel channel were quantified by digital image correlation (DIC) using fluorescent microspheres embedded in the fibrin gel during actuation at vacuum pressures ranging from 20 to 80 kPa (**Fig. 3D**). As expected, peak y strain was positive and increased with increasing vacuum pressure (**Fig. 3E**). At -80 kPa, the mean peak strain magnitude reached 5.92% in the 200-µm-wide channel, 2.93% in the 500-µm-wide channel, and 1.12% in the 1000-µm-wide channel (**Fig. 3E**). By contrast, peak strain magnitudes along the gel channel were lower, remaining near zero in the 200-µm-wide channel and reaching mean values of 0.86% and 1.03% in the 500- and 1000-µm-wide channels, respectively (**Fig. 3E**). The design with the 1000-µm-wide channel was excluded from further experiments due to the insufficient strain magnitudes under actuation. While the 200-µm-wide channel offered the widest range of y strain magnitudes, preliminary experiments revealed that it did not support the formation of microvascular networks. Instead, hPAECs formed a single large vessel with little surrounding extracellular matrix available for tumor cell extravasation (**Supplementary Fig. S1**). The 500-µm-wide channel design and a vacuum pressure of 80 kPa were therefore selected for further experiments, as these parameters produced physiological y strains of approximately 3% [4, 5].

To independently validate the x- and y-strain magnitudes measured by DIC and further estimate the out-of-plane z-strain, we performed finite element analysis (FEA) of the 500-μm-wide gel channel actuated at a vacuum pressure of 80 kPa (**Fig. 3F**). The model predicted average x- and y-strain magnitudes of 0.43% and 3.21%, respectively, in the actuated region of the gel (**Fig. 3F**), in close agreement with the corresponding DIC measurements of 0.86% and 2.93% (**Fig. 3E**). The model further predicted an average z-strain of −7.23% (**Fig. 3F**). Together, these complementary experimental and computational approaches demonstrate that the mechanically actuated device reproduces the multidirectional strains, extensional in the x-y directions and compressive in the z direction, consistent with those experienced by lung capillaries during normal breathing (**Fig. 3A**).

### Mechanically actuated lung-specific vessels undergo a physiological reduction in diameter

After selecting the final design of the mechanically actuated device and characterizing the strain magnitudes within the gel, we next examined device performance in the presence of microvascular networks. Lung-specific microvascular networks self-assembled over 7 days without mechanical actuation but in the presence of transient flow driven by a hydrostatic pressure gradient, resulting in vessels aligned along the gel channel (**Fig. 4A**). Perfusion of fluorescent dextran (70 kDa) further confirmed that the vessels were fully perfusable (**Fig. 4B**). Formed microvascular networks were then subjected to cyclic actuation at a vacuum pressure of 80 kPa and a frequency of 0.2 Hz, corresponding to a physiological breathing rate [7], and imaged before and during actuation. Analysis of vessel diameters in the x-y plane revealed a reduction in vessel diameter that was consistent across devices, ranging from −1.99% to −13.08%, with a mean reduction of 6.02% (**Fig. 4C**). Because vessel diameters varied within the microvascular networks and smaller vessels appeared to undergo greater proportional narrowing, we next investigated whether actuation-induced vessel deformation depended on vessel size. Paired vessel diameters were analyzed using a log-transformed difference-versus-mean approach adapted from Bland and Altman [31] and Cole [32]. Relative diameter change was expressed as the natural logarithm of the actuated-to-not actuated diameter ratio and plotted against the mean natural logarithm of the paired vessel diameters (**Fig. 4D**). Linear regression revealed a significant relationship between vessel size and relative diameter change (R² = 0.385, P = 0.0001), indicating that smaller vessels underwent greater relative reductions in diameter than larger vessels (**Fig. 4D**). Consistent with the experimental observations, FEA of a single vessel embedded in fibrin within a 500-μm-wide channel qualitatively reproduced the reduction in vessel diameter during actuation at a vacuum pressure of 80 kPa (**Fig. 4E**). Because the vessel was modeled as an empty lumen without accounting for fluid resistance, the model was used to assess the direction of vessel deformation rather than its magnitude. Overall, these results show that the mechanically actuated device recapitulates the physiological reduction in vessel diameter experienced by lung capillaries during breathing [30].

**Figure 4.**
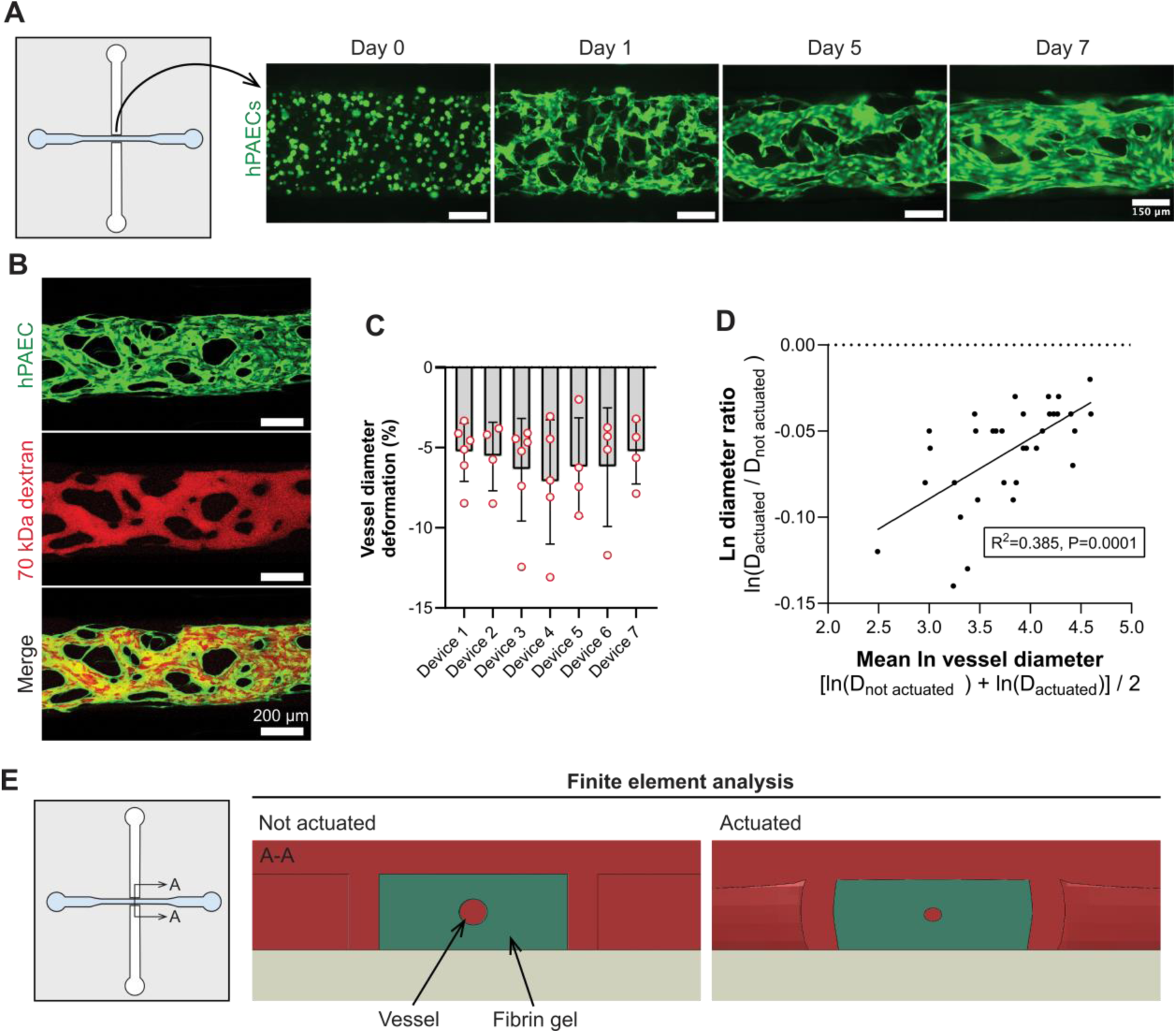
Mechanically actuated lung-specific vessels undergo a physiological reduction in diameter. (A) Representative epifluorescence microscopy images of microvascular network formation over 7 days in the actuation region of the mechanically actuated device. Scale bar is 150 μm. (B) Representative confocal microscopy images of dextran-perfused microvascular networks showing full vessel perfusability. Scale bar is 200 μm. (C) Characterization of vessel diameter deformation under actuation. n = 4-6 vessels per device from 7 devices. (D) Linear regression of the natural logarithm of the actuated-to-not actuated vessel diameter ratio against the mean natural logarithm of the actuated and not actuated vessel diameters. n = 33 vessels from 7 devices. (E) Cross-sectional views of a finite element model of a single vessel embedded in fibrin within a 500-μm-wide channel, shown in the not actuated state and during actuation at a vacuum pressure of 80 kPa.

### Mechanical actuation preserves lung-specific vessel viability, permeability, and morphology

As our device applies breathing-like physiological mechanical actuation, it was important to verify that it did not compromise the viability, structure, and function of the embedded microvascular networks. Lung-specific microvascular networks were cultured for 24 h with or without cyclic actuation at a vacuum pressure of 80 kPa and a frequency of 0.2 Hz. Cell viability was assessed by staining samples with NucBlue Live and propidium iodide before or after the 24-hour period (**Fig. 5A**). Quantitative analysis revealed that high cell viability (> 90%) was maintained across conditions and there were no statistically significant differences between groups (**Fig. 5B**). Similarly, mechanical actuation did not significantly affect vascular permeability (**Fig. 5C**) or morphological traits (**Fig. 5D-H**). These results indicate that subsequent functional assays, including tumor cell extravasation, are unlikely to be confounded by actuation-induced defects in vascular viability, permeability, or morphology.

**Figure 5.**
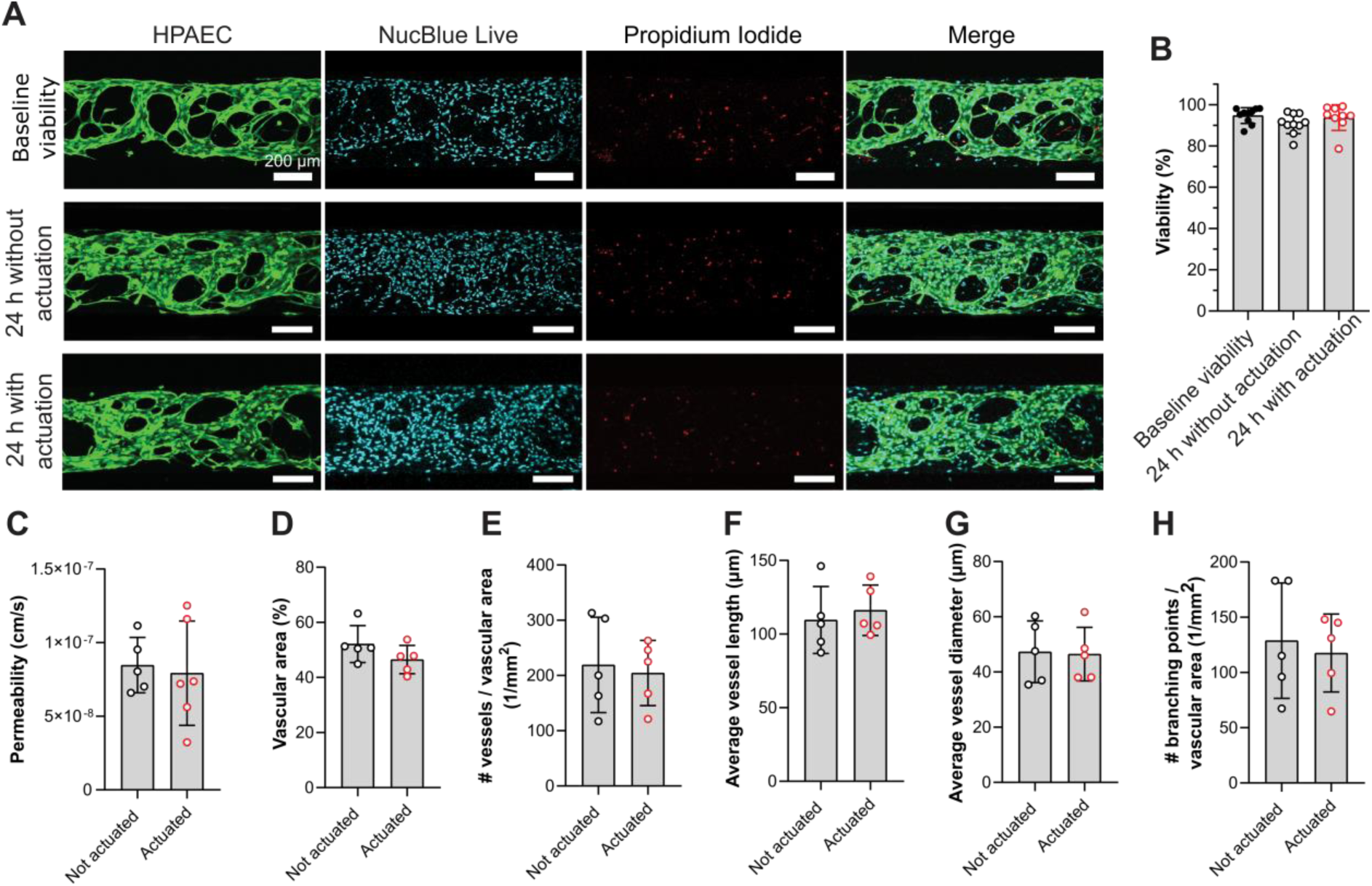
Mechanical actuation preserves lung-specific vessel viability, permeability, and morphology. (A) Representative confocal microscopy images of hPAEC microvascular networks on day 5 (baseline viability), after 24 h without actuation, and after 24 h with actuation. Samples were stained with NucBlue Live and propidium iodide to label all the nuclei and the nuclei of dead cells, respectively. Scale bar is 200 μm. (B) Quantification of cell viability in hPAEC microvascular networks on day 5 (baseline viability), after 24 h without actuation, and after 24 h with actuation. n = 9-10 devices. (C) Permeability analysis of hPAEC microvascular networks that were either not actuated or actuated for 24 h. n = 5-6 devices. (D-H) Morphological analysis of hPAEC microvascular networks that were either not actuated or actuated for 24 h, including (D) projected vessel area fraction, (E) vessel density, (F) average vessel length, (G) average vessel diameter, and (H) branching point density. n = 5 devices.

### Co-actuation of lung-specific microvascular networks and perfused tumor cells increases tumor cell extravasation

In a final step, we used the mechanically actuated device to investigate the effect of breathing-like mechanical actuation on tumor cell extravasation. We sought to distinguish the effects of mechanical actuation on the vasculature from those occurring during tumor cell extravasation by conducting a “pre-actuation” and a “co-actuation” study, respectively. In the pre-actuation study, lung-specific microvascular networks grown over 5 days were cultured for an additional 24 h with or without cyclic actuation (-80 kPa, 0.2 Hz). MDA-MB-231 cells were then perfused and allowed to extravasate under static conditions (**Fig. 6A**). This approach allowed us to determine whether prior mechanical actuation of the vasculature alone altered subsequent tumor cell extravasation. In contrast, in the co-actuation study, the 24-hour period with or without actuation began immediately after the MDA-MB-231 cells were perfused and continued while the tumor cells extravasated (**Fig. 6A**). In this scenario, mechanical actuation is applied to both the vasculature and the tumor cells. In both studies, lung-specific microvascular networks perfused with tumor cells were live-imaged on day 7 by confocal microscopy to quantify tumor cell extravasation rates (**Fig. 6B**).

**Figure 6.**
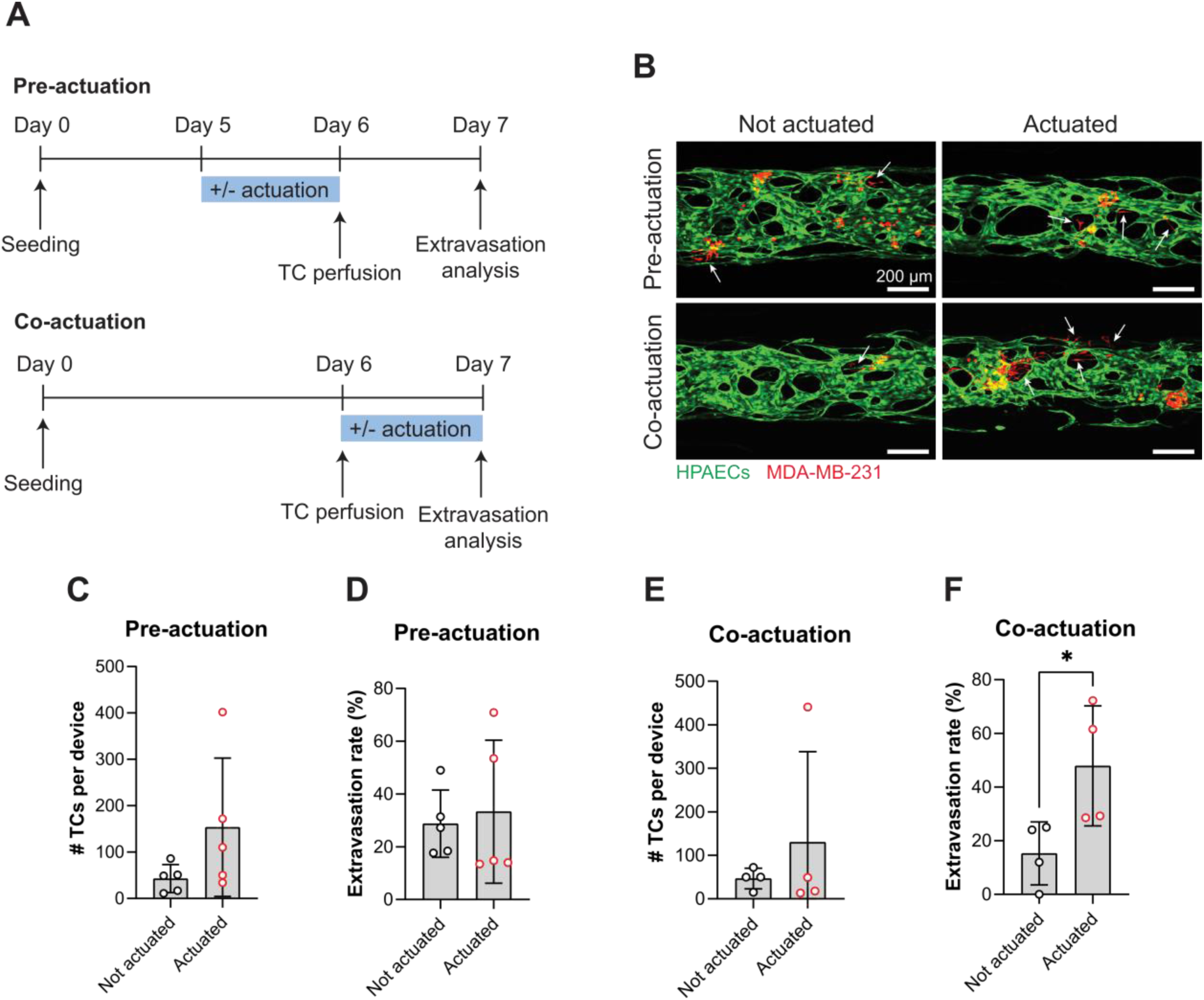
Co-actuation of lung-specific microvascular networks and perfused tumor cells increases tumor cell extravasation. (A) Experimental timeline for the pre-actuation and co-actuation studies. In the pre-actuation study, lung-specific microvascular networks were cultured with or without breathing-like cyclic actuation for 24 h before tumor cell (TC) perfusion. Tumor cells were then perfused and allowed to extravasate under static conditions. In the co-actuation study, tumor cells were perfused at the beginning of the 24-h actuation period and allowed to extravasate while the microvascular networks were subjected to breathing-like cyclic actuation or maintained under static conditions. (B) Representative confocal microscopy images of hPAEC microvascular networks and extravasated tumor cells (white arrows) on day 7 under the four experimental conditions. Scale bar is 200 μm. (C-D) Tumor cell extravasation analysis of the pre-actuation study, showing (C) total number of tumor cells (TCs) per device and (D) tumor cell extravasation rate in the not actuated and actuated conditions. n = 5 devices. (E-F) Tumor cell extravasation analysis of the co-actuation study, showing (E) total number of tumor cells (TCs) per device and (F) tumor cell extravasation rate in the not actuated and actuated conditions. n = 4 devices. * indicates p < 0.05.

In the pre-actuation study, the total number of tumor cells per device was similar between the not actuated and actuated conditions (**Fig. 6C**). Similarly, there was no significant difference in the tumor cell extravasation rate between the two groups (**Fig. 6D**). The total number of tumor cells per device was also similar between conditions in the co-actuation study (**Fig. 6E**), whereas the tumor cell extravasation rate was considerably higher in the actuated condition relative to the not actuated condition (47.89% versus 15.25%) (**Fig. 6F**). Overall, these results show that prior actuation of the vasculature alone does not alter subsequent tumor cell extravasation, whereas mechanical actuation during the extravasation process promotes tumor cell extravasation. As such, we propose that breathing-like mechanical actuation is an important factor to include and consider when studying cancer metastasis.

## DISCUSSION

The goal of this study was to develop a mechanically actuated lung-specific microvascular model that mimics breathing mechanics and to determine how endothelial cell type and breathing-like actuation affect tumor cell extravasation. First, we characterized lung-specific microvascular networks in terms of barrier function and structure. The permeability of lung-specific microvascular networks to 70-kDa dextran (∼6 x 10^−8^ cm/s) was of the same order of magnitude as the reported permeability of lung microvasculature to albumin in sheep (∼10^−8^ cm/s) [33], despite differences in tracer properties and measurement approaches. Notably, this permeability was substantially lower than values reported for human lung microvascular endothelial monolayers exposed to 70-kDa dextran (∼10^−5^ cm/s) [34], suggesting that the self-assembled three-dimensional microvascular networks reproduce a relatively tight physiological vascular barrier. We also showed that, compared to traditional iHUVEC microvascular networks, lung-specific networks were denser and composed of shorter vessels. This morphology is qualitatively consistent with the alveolar capillary bed, which forms a dense, highly interconnected network of short capillary segments to maximize gas exchange [5, 35, 36].

When iHUVEC and lung-specific microvascular networks were perfused with MDA-MB-231 cells, we observed a marked increase in tumor cell extravasation in the lung-specific vasculature despite similar total numbers of tumor cells per device. This suggests that the higher extravasation rate was not driven by greater overall tumor cell retention, but rather by more efficient transmigration across the lung-specific vasculature. This finding is consistent with previous work showing that MDA-MB-231 cells can harbor gene programs that promote breast cancer metastasis to the lung [11]. Importantly, our results further demonstrate that changing the endothelial cell source alone can alter tumor cell extravasation, highlighting the value of organ-specific vasculature in models of metastatic dissemination.

In the next step, we aimed to recapitulate not only lung-specific vasculature but also the micromechanical environment of the alveolus during normal breathing. We used two independent approaches, DIC and FEA, to characterize multidirectional strains within the fibrin matrix of the mechanically actuated device. The selected device and actuation parameters (500-μm-wide gel channel and 80 kPa vacuum pressure) generated tensile strains of approximately 0.5-1% and 3% in the x and y directions, and a compressive strain of -7% in the z direction, with good agreement between DIC and FEA. Reported estimates of alveolar deformation vary depending on the experimental model, lung-volume range, and measurement approach [6], and are often derived from changes in alveolar geometry or surface area rather than multidirectional strain measurements. Nevertheless, the strains generated by our device are comparable to reported estimates of approximately 4-10% for linear alveolar deformation during tidal breathing [1–6].

Importantly, deformation of the surrounding matrix translated into cyclic deformation of the embedded lung-specific microvasculature, with vessel diameters decreasing by 2-13% and by approximately 6% on average. A reduction in vessel diameter was also qualitatively reproduced by FEA, consistent with animal studies showing that increasing lung inflation flattens and compresses alveolar capillaries [37–39]. More recently, Regan et al. directly measured capillary deformation in intact mouse lungs and reported an approximately 3% reduction in capillary diameter under negative-pressure ventilation mimicking spontaneous breathing [30]. Although the experimental conditions are not directly equivalent, the magnitude of vessel narrowing observed in our model is comparable to these direct measurements in the lung. Moreover, we found that smaller vessels exhibited greater relative diameter reductions. This size dependence may be relevant to the design and comparison of engineered vascular models, as differences in initial vessel diameter may contribute to differences in relative deformation under mechanical actuation. Together, these findings support the physiological relevance of the mechanical response of the lung-specific microvasculature in our model.

Following mechanical characterization of the device and quantification of vessel deformation, we next assessed whether prolonged breathing-like cyclic actuation for 24 h at 0.2 Hz affected the viability, permeability, or morphology of the lung-specific microvascular networks. Cell viability remained above 90% across conditions, indicating that 24 h of mechanical actuation did not compromise vessel viability. This finding is consistent with previous studies showing that lung endothelial cells tolerate low-magnitude cyclic strain. In hPAEC monolayers, Birukov et al. found only a small increase in cell death after 6 h of 5% cyclic strain, while 18% strain induced greater cell death [40]. Importantly, cell death did not further increase between 6 and 24 h [40]. Previous studies have also demonstrated magnitude-dependent effects of cyclic strain on lung endothelial barrier function, with 5% strain promoting barrier recovery or reducing susceptibility to agonist-induced barrier disruption compared with 18% strain [40–42]. Similarly, in the lung-on-a-chip developed by Huh and colleagues, alveolar-capillary barrier permeability remained unchanged during 4 h of cyclic actuation at strain levels of 5-15% [18]. These findings are consistent with the preserved vascular permeability observed in our mechanically actuated networks. A barrier-preserving effect of cyclic stretch has also been demonstrated in three-dimensional vessels. Zeinali et al. showed that 24 h of cyclic stretch at 0.2 Hz significantly reduced 70-kDa dextran permeability in perfusable endothelialized single vessels [22]. Mechanical actuation also did not alter the morphological characteristics of our formed microvascular networks. In contrast, Ferrari et al. reported increased vascular density, branch number, total branch length, and junction number after 24 h of cyclic stretch in an intervessel network undergoing vasculogenesis [23], suggesting that mechanical stimulation may have stronger effects on vascular structure during network formation. Consistent with the stability observed in our formed networks, Paek et al. recently found no significant vascular remodeling in self-assembled microvasculature within normal airway constructs subjected to repeated bronchoconstriction-like compression, whereas substantial remodeling occurred in corresponding asthmatic constructs [29]. Together, our results demonstrate that the viability, barrier function, and overall architecture of lung-specific microvascular networks are maintained during 24 h of breathing-like mechanical actuation.

In the final step of this study, we tested the effect of breathing-like mechanical actuation on MDA-MB-231 tumor cell extravasation. We found that pre-actuation of the lung-specific microvascular networks followed by tumor cell perfusion and extravasation under static conditions did not influence tumor cell extravasation. This finding is consistent with our earlier results showing that physiological mechanical actuation maintains vascular barrier integrity and further suggests that prior actuation does not induce persistent endothelial changes that facilitate subsequent tumor cell extravasation. In contrast, when lung-specific microvascular networks and tumor cells were co-actuated during extravasation, we observed a marked increase in tumor cell extravasation compared with the not actuated condition. These data suggest that mechanical actuation promotes tumor cell extravasation only when applied during tumor-endothelial interaction. To our knowledge, no previous studies have investigated the effect of physiological breathing-like mechanical actuation on tumor cell extravasation in lung vasculature. Nevertheless, several studies have shown that applied mechanical stimulation, including cyclic strain, can alter tumor cell phenotype and promote invasive behavior [43–46]. Cyclic strain has also been shown to enhance tumor cell adhesion to endothelial cells and alter metastatic behavior [47]. In the context of tumors established in the lung, López-Alonso and colleagues further showed that mechanical ventilation of tumor-bearing mice increased metastatic dissemination and linked this effect to increased tumor cell invasiveness observed following cyclic strain in vitro [16]. More directly relevant to the lung as a metastatic organ, Huang and colleagues reported that mechanical ventilation increased the retention of intravenously injected 4T1 breast cancer cells in the lung microvasculature and, in a separate spontaneous breast cancer metastasis model, increased postoperative lung metastatic burden [15]. However, mechanical ventilation in their model induced lung injury, increased vascular permeability, and disrupted lung microvascular integrity [15], providing a substantially different vascular environment from the physiological actuation used here. Intravital imaging studies have also shown that metastatic tumor cells become physically lodged within lung capillaries, although the imaged lung region was immobilized using a vacuum-stabilized window, preventing the study of tumor cell behavior under local breathing-induced deformation [13]. More recently, using the Crystal Ribcage platform, Banerji and colleagues demonstrated that metastatic breast tumor cells confined within lung capillaries deform in response to changes in alveolar pressure, directly showing that respiratory mechanics are transmitted to tumor cells within the lung microvasculature [17]. However, tumor cell extravasation was not observed in this model, which may in part reflect the technical challenge of maintaining the ex vivo preparation over the longer timescales required to observe extravasation. Our mechanically actuated lung microvascular model addresses this gap by combining physiological breathing-like deformation with perfusable lung-specific microvascular networks, enabling the direct study of tumor cell extravasation under dynamic mechanical conditions over prolonged periods.

In conclusion, we developed a microphysiological system incorporating lung-specific vasculature and a single-layer mechanical actuation platform capable of reproducing breathing mechanics. The system enables direct visualization and quantitative assessment of extracellular matrix and vascular deformations that were consistent with reported physiological ranges. Mechanical actuation preserved vessel viability, structure, and barrier function, making the system well suited for studying tumor cell extravasation under dynamic mechanical conditions. Using this system, we showed that mechanical actuation during tumor cell extravasation increased extravasation, whereas prior actuation alone did not affect subsequent extravasation. This finding highlights breathing mechanics as a previously underexplored component of the metastatic lung microenvironment and suggests that organ-specific mechanical activity should be considered when modeling dynamic tissue microenvironments. More broadly, the platform could be adapted to reproduce other forms of cyclic tissue deformation, including cardiac contraction, muscle activation, and the mechanical activity of organs such as the gut, bladder, and intervertebral disc [48], enabling the study of a range of physiological and pathological processes.

## METHODS

### Cell culture

Human umbilical vein endothelial cells (HUVECs; Lonza, C2519A) were immortalized and transduced to stably express cytoplasmic blue fluorescent protein as previously described [49]. Immortalized HUVECs (iHUVECs) at passage 9 were cultured in 75 cm² flasks in VascuLife VEGF Endothelial Medium containing all supplements provided in the kit, except heparin sulfate, which was added at one quarter of the recommended concentration (Lifeline Cell Technology, LL-0003). Green fluorescent protein-expressing human pulmonary artery endothelial cells (hPAECs; Angio-Proteomie, cAP-0120GFP) at passage 7 were cultured in 75 cm² flasks pre-coated for 5 min with Quick Coating Solution (Angio-Proteomie, cAP-01), using the same VascuLife medium as for iHUVECs supplemented with an additional 10% fetal bovine serum (FBS; Thermo Fisher Scientific, A5670701). Primary human normal lung fibroblasts (hNLFBs; LifeLine Cell Technology, FC-0049) at passage 6 were cultured in 75 cm² flasks in FibroLife S2 Fibroblast Medium containing all supplements provided in the kit (LifeLine Cell Technology, LL-0011). MDA-MB-231 cells (ATCC, HTB-26) stably expressing cytoplasmic tdTomato were cultured in in 75 cm² flasks in Dulbecco’s Modified Eagle Medium (Thermo Fisher Scientific, 11995065) supplemented with 10% FBS (Thermo Fisher Scientific, A5670701) and 1% Penicillin-Streptomycin (Thermo Fisher Scientific, 15140122). All cells were maintained at 37 °C in a humidified incubator with 5% CO₂.

### Design and fabrication of static microfluidic device

Static microfluidic devices [50] were designed and fabricated as previously described [51, 52]. Molds were designed in AutoCAD (Autodesk), imported into Fusion 360 (Autodesk) to generate the corresponding toolpaths, and fabricated using a micro-CNC milling machine (Bantam Tools). Polydimethylsiloxane (PDMS; Dow Corning Sylgard 184, Ellsworth Adhesives) was mixed at a 10:1 w/w ratio of base to crosslinker, degassed for 30 min, poured into the molds, degassed for an additional 30 min, and cured overnight at 60 °C. Individual devices were cut, and biopsy punches (Integra Miltex) were used to punch ports into the devices. The devices were then sterilized in an autoclave, and glass coverslips (No. 1, VWR) were sonicated in pure ethanol and dried. The devices and coverslips were plasma-treated using a plasma cleaner (Harrick Plasma) and bonded together. The bonded devices were placed in a 70 °C oven for 48 h to restore hydrophobicity.

### Formation of microvascular networks in static microfluidic device

After detachment from cell culture flasks with Accutase (Millipore, SCR005), hNLFBs were resuspended and mixed either with iHUVECs or hPAECs in cold VascuLife medium containing 3.6 U/mL thrombin (Sigma-Aldrich, T4648). Each cell mixture was combined with an equal volume of 5 mg/mL fibrinogen (Sigma-Aldrich, F8630) and immediately injected into the central gel channel of the static microfluidic device. The final cell densities for the iHUVEC-hNLFB co-culture were 9 x 10^6^ iHUVECs/mL and 1.5 x 10^6^ hNLFBs/mL, and the final cell densities for the hPAEC-hNLFB co-culture were 9 x 10^6^ hPAECs/mL and 2 x 10^6^ hNLFBs/mL. Devices were placed in a humidified incubator at 37 °C and 5% CO₂ for 10 min for gel polymerization, after which pre-warmed medium was added to the device side channels. As described for cell culture in flasks, VascuLife medium was used for the iHUVEC-hNLFB co-culture, while VascuLife supplemented with 10% FBS was used for the hPAEC-hNLFB co-culture. The remaining endothelial cells were replated in 75 cm² flasks to be used 3 days later to seed cell monolayers on the vertical sides of the gels in the devices. Medium was changed daily, and on day 2 of culture, 1 mL syringes (plungers removed and barrels shortened) were placed in the ports of one of the media channels of each device. Each syringe was filled with 200 μL of medium to establish a hydrostatic pressure gradient across the gel channel and generate interstitial flow, the direction of which was alternated daily.

On day 3, cell monolayers were seeded on the vertical sides of the gels in the devices. Syringes were removed, medium was aspirated from both side channels, and the channels were filled with a 30 μg/mL fibronectin (Millipore Sigma, FC010) solution and incubated at 37 °C and 5% CO₂ for 30 min. Replated iHUVECs and hPAECs were detached with Accutase and resuspended at 1 x 10^6^ cells/mL in their respective media. The fibronectin solution was aspirated and 40 μL of the appropriate endothelial cell suspension, equivalent to 40,000 cells, was added to one of the medium channels. The devices were tilted at an angle of 90° for 5 min so that the endothelial cells could attach to the vertical gel surface. These steps were repeated for the second medium channel. The devices were cultured without flow for 24 h to allow the endothelial cells to firmly adhere to the gel, after which the syringes were replaced and interstitial flow was reestablished. Medium was changed daily until day 7.

### Vascular permeability analysis in static microfluidic device

On day 7, vascular permeability was analyzed as previously described [52, 53]. Briefly, pre-warmed 0.1 mg/mL Texas Red 70-kDa dextran (Thermo Fisher Scientific, D1864) was added to each medium channel of the static microfluidic device. Devices were immediately imaged at 37 °C on a confocal microscope (Olympus, FV1000) using a 10x objective. Two sets of z-stack images were acquired with a z-step of 5 μm and an x-y resolution of 640 x 640 pixels at 0 and 12 min. Vascular permeability was quantified by analyzing the changes in the fluorescence intensity within the vessels and the surrounding extracellular matrix over time, as previously described [52].

### Vascular morphology analysis in static microfluidic device

Z-stack images of dextran-perfused vessels acquired for vascular permeability analysis were used to analyze vascular morphology in ImageJ (NIH), as previously described [54, 55]. Maximum-intensity projections were generated and thresholded using the “Huang” method. Images were despeckled, and dark and bright outliers were removed using the “Remove Outliers” function with radii of 5 and 2 pixels and a threshold of 50. The images were then skeletonized, and morphological features were extracted using the “Analyze Skeleton” function. The average vessel diameter, *D*, was calculated as: 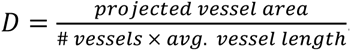.

### Tumor cell extravasation assay in static microfluidic device

On day 7 of microvascular network culture, MDA-MB-231 cells were detached with 2 mM ethylenediaminetetraacetic acid (EDTA) (Gibco, 15575-038) and resuspended in VascuLife medium supplemented with 10% FBS at a concentration of 1 x 10^6^ cells/mL. One of the medium channels of the static microfluidic device was emptied and 70 μL of cell suspension, equivalent to 70,000 cells, was introduced into the channel. Medium was changed daily with VascuLife medium supplemented with 10% FBS. Devices were live-imaged 42 h after tumor cell perfusion on a confocal microscope (Olympus, FV1000) using a 10x objective. Z-stack images were acquired with a z-step of 5 μm and an x-y resolution of 640 x 640 pixels. Maximum-intensity projections and orthogonal views were used in Imaris software (BitPlane) to manually count intravascular and extravasated cells. Tumor cells in the process of extravasating or fully extravasated were counted as extravasated. The total number of cells per device was calculated by adding all the cells in the gel (intravascular and extravasated). The extravasation rate was calculated as: 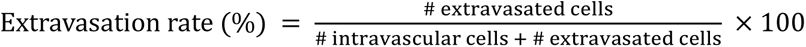

### Design and fabrication of mechanically actuated microfluidic device

The molds for the mechanically actuated microfluidic devices were designed and fabricated similarly to those of the static microfluidic device. In addition to being poured into molds, PDMS was also poured into a Petri dish to fabricate donut-shaped syringe supports. After overnight curing at 60 °C, syringe supports (9 mm outer diameter, 4 mm inner diameter) were punched out of the PDMS in the Petri dish and autoclaved. Individual mechanically actuated PDMS devices (cut, punched, and autoclaved) were bonded to clean glass coverslips (No. 1, VWR) with a partially cured spin-coated layer of PDMS. For the spin-coating procedure, a PDMS mixture (10:1 w/w ratio of base to curing agent) was prepared and degassed under vacuum. For each device, a glass coverslip was spin-coated with 1 mL of the PDMS mixture using a spin coater (Laurell, WS-400B-6NPP) at 2480 rpm for 5 min, generating a ∼15 μm thick PDMS layer. The spin-coated coverslip was partially cured at 70 °C for 10 min. A PDMS device was then preheated at 70 °C, carefully aligned, and placed onto the spin-coated coverslip. This procedure was performed individually for each device. The assembled devices were cured overnight at 70 °C to establish irreversible bonding between the PDMS device and the spin-coated glass substrate. The following day, the syringe supports were plasma-bonded on top of the devices. Devices were placed in a 70 °C oven for 48 h to restore hydrophobicity.

### Pneumatic actuation system and mechanical stimulation

A custom pneumatic vacuum actuation system was developed to control the magnitude and frequency of the applied negative pressure. A centralized laboratory vacuum source served as the primary vacuum supply and was connected to a precision vacuum regulator (Grainger, IRV10-N07BG) to control and monitor the applied pressure. The regulated vacuum was directed through a solenoid valve (SMC Pneumatics, S070C-6CG-32) to an eight-port manifold (Pawfly, AFC834). The outlet ports of the manifold were connected to the pneumatic channels of the microfluidic devices using silicone tubing (Vanguard, FGVP51135-11-25), allowing simultaneous actuation of multiple devices. The solenoid valve was controlled by a pulse-width modulation signal generator (Pemenol, B07W229NVP) to regulate the actuation frequency. Unless otherwise specified, devices were mechanically actuated at −80 kPa and 0.2 Hz, corresponding to a physiological human breathing rate at rest [7].

### Digital image correlation analysis

Mechanically actuated devices were fabricated with three different gel channel widths: 200 μm, 500 μm, and 1000 μm. Yellow-green, 1.0 μm fluorescent polystyrene microspheres (Thermo Fisher Scientific, F13081) were suspended at a concentration of 1 x 10^7^ beads/mL in a 3.6 U/mL thrombin solution. The thrombin solution was mixed with a 5 mg/mL fibrinogen (Sigma-Aldrich, F8630) solution at a 1:1 ratio and immediately injected into the gel channel of a mechanically actuated device. Devices were placed in a humidified incubator at 37 °C and 5% CO₂ for 15 min for gel polymerization, after which VascuLife medium was added to the device medium ports. The next day, the devices were connected to the pneumatic actuation system and mechanically actuated at −80 kPa and 0.2 Hz. An epifluorescence microscope (Nikon Eclipse Ti-U Inverted Fluorescence Microscope) was used to acquire videos of the actuated device at 100 frames per second using NIS-Elements BR software. Digital image correlation (DIC) was performed on the images extracted from each video using a Python framework (Python 3.14.6, μDIC package) [56]. Strains were evaluated in the actuated region of the device over an area with a length of 1200 μm (along the channel, x strain) and half the gel channel width (across the channel, y strain) using a grid with a patch size of 120 μm along the channel and 50 μm across the channel. The non-actuated reference state was identified as the frame with the minimum y strain within an actuation cycle. Peak x and y strains were extracted relative to that reference image. For each device, strains were averaged across all patches of the grid and across three load cycles.

### Finite element analysis of mechanically actuated device deformation

Finite element analysis (FEA) was used to independently predict the mechanics of the mechanically actuated device. Models were solved with Abaqus using a quasi static analysis (Abaqus 2025, Dassault Systèmes). The PDMS and gel were modeled with linear hexahedral hybrid elements (C3D8H) in combination with hyperelastic Neo-Hookean material models (PDMS [57]: C10 = 237.5 kPa, D1 = 4.23 x 10^-5^ kPa^-1^, 94604 nodes, 86911 elements, Gel [58]: C10 = 8.36 x 10^-3^ kPa, D1 = 1.2 kPa^-1^, 29295 nodes, 24640 elements) using a hybrid formulation. The glass was modeled with linear hexahedral elements (C3D8R) and a linear elastic material (E = 7.0 x 10^7^ kPa, ν = 0.2 [59], 12051 nodes, 7932 elements). The influence of the discretization was considered converged when strain values changed by less than 5% compared to a mesh with double the edge length. The bottom surface of the glass coverslip was fixed. Contact between the PDMS and glass was modeled with a tied constraint. The outer surface of the gel was constrained to the surface of the PDMS and glass within the channel. Device actuation was modeled by applying a pressure of -80 kPa to all surfaces within the vacuum channel. Strains along the channel (x strain), across the channel (y strain) and through the depth of the channel (z strain) were averaged through the thickness of the gel within the actuated region corresponding to the area used for DIC analysis.

### Formation of lung-specific microvascular networks in mechanically actuated microfluidic device

After detachment from cell culture flasks with Accutase (Millipore, SCR005), hNLFBs and hPAECs were resuspended in cold VascuLife medium supplemented with 10% FBS and containing 3.6 U/mL thrombin (Sigma-Aldrich, T4648). The hNLFB and hPAEC suspensions were mixed, and the resulting cell mixture was combined with an equal volume of 5 mg/mL fibrinogen (Sigma-Aldrich, F8630) and immediately injected into the gel channel of the mechanically actuated device. The final cell densities were 5 x 10^6^ hPAECs/mL and 1.5 x 10^6^ hNLFBs/mL. Devices were placed in a humidified incubator at 37 °C and 5% CO₂ for 15 min for gel polymerization, after which 3 mL syringes were placed in the syringe supports, and VascuLife medium supplemented with 10% FBS was added to create a 1 cm H_2_O hydrostatic pressure gradient along the gel channel. Medium was changed daily to maintain the hydrostatic pressure gradient.

### Quantification of vessel deformation during mechanical actuation

Lung-specific microvascular networks were mechanically actuated at −80 kPa and 0.2 Hz and imaged using an epifluorescence microscope (Nikon Eclipse Ti-U Inverted Fluorescence Microscope). Videos were acquired at 100 frames per second using NIS-Elements BR software. Representative frames corresponding to the not actuated and maximally actuated states were extracted from the videos and analyzed in Fiji. For each device, the diameters of 4-6 vessels were manually measured at the same location in the not actuated and actuated frames. The relative change in vessel diameter was calculated as: (*D*_actuated_ − *D*_not actuated_)/*D*_not actuated_ × 100.

To determine whether proportional vessel narrowing varied with vessel size, paired vessel diameters were analyzed using a log-transformed difference-versus-mean approach adapted from Bland and Altman [31]. For each vessel, proportional diameter change was expressed as the natural-log diameter ratio, ln(*D*_actuated_/*D*_not actuated_), and vessel size was represented by the mean of the paired log-transformed diameters, [ln(*D*_not actuated_) + ln(*D*_actuated_)]/2. The natural-log ratio provides a symmetric measure of fractional change and, when multiplied by 100, can be interpreted as the symmetric percentage difference described by Cole [32]. A simple linear regression was used to test the association between proportional diameter change and vessel size. Device-specific slopes and intercepts were first compared. Because no significant differences were detected, the pooled regression across vessels was reported.

### Finite element analysis of vessel deformation

A second FEA modeled the gel with a cylindrical hole to represent a vessel and predict changes in vessel diameter. The vessel was modeled as an empty cylinder with an internal pressure of 49 Pa due to the hydrostatic pressure gradient generated by the syringes. This approach did not explicitly model the fluid within the vessel, allowing for a simplified analysis. Therefore, the vessel dimensions were only qualitatively compared before actuation and at −80 kPa in both the y direction and the z direction.

### Cell viability analysis following mechanical actuation

Cell viability was assessed using the Blue/Red ReadyProbes Cell Viability Imaging Kit (Thermo Fisher Scientific, R37610). Lung-specific microvascular networks were cultured for 24 h with or without cyclic mechanical actuation at -80 kPa and 0.2 Hz. Viability was measured on day 5 (baseline), and after the 24-h period. Microvascular networks were incubated with NucBlue Live (2 drops/mL) and propidium iodide (2 drops/mL) for 30 min in a humidified incubator at 37 °C and 5% CO₂. Devices were immediately imaged on a confocal microscope (Olympus, FV1000) using a 10x objective. Z-stack images were acquired with a z-step of 5 μm and an x-y resolution of 800 x 800 pixels. NucBlue and propidium iodide channels were analyzed separately in ImageJ (NIH). Each z-stack was thresholded and the resulting binary images were processed using the “Despeckle” and “Watershed” functions. NucBlue-positive nuclei and propidium iodide-positive nuclei were then quantified using the “3D Objects Counter” plugin with channel-specific object-size thresholds. Viability was calculated as: 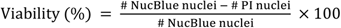.

### Vascular permeability and morphology analyses following mechanical actuation

Lung-specific microvascular networks were cultured for 24 h with or without cyclic mechanical actuation at -80 kPa and 0.2 Hz. At the end of the 24-h period, vascular permeability and morphology were analyzed as described above.

### Tumor cell extravasation assay under mechanical actuation

In the pre-actuation study, lung-specific microvascular networks were cultured with or without breathing-like cyclic actuation at -80 kPa and 0.2 Hz for 24 h, starting on day 5, before tumor cell perfusion on day 6. Tumor cells were then perfused and allowed to extravasate under static conditions. In the co-actuation study, tumor cells were perfused at the beginning of the 24-h actuation period on day 6 and allowed to extravasate while the microvascular networks were subjected to breathing-like cyclic actuation at -80 kPa and 0.2 Hz or maintained under static conditions. MDA-MB-231 cells were detached with 2 mM ethylenediaminetetraacetic acid (EDTA) (Gibco, 15575-038) and resuspended in VascuLife medium supplemented with 10% FBS at a concentration of 5 x 10^5^ cells/mL. The gel channel ports of the mechanically actuated device were emptied and 100 μL of cell suspension, equivalent to 50,000 cells, was introduced into one of the ports. Medium was changed daily with VascuLife medium supplemented with 10% FBS. Devices were live-imaged 24 h after tumor cell perfusion, on day 7, as described above, and images were analyzed as described above to calculate the tumor cell extravasation rate.

### Statistical analysis

Data in bar graphs were presented as mean ± standard deviation. Individual data points represent single microfluidic devices. Statistical analysis was conducted using GraphPad Prism 11. Data were assessed for normality using the normality tests available in GraphPad Prism, as permitted by sample size. For comparisons between two independent conditions (iHUVECs versus hPAECs or not actuated versus actuated), an unpaired two-tailed Student’s t-test was used for normally distributed data. Welch’s t-test was used when significant differences in variance were detected. For non-normally distributed data, a Mann-Whitney test was performed. When comparing the three independent conditions in the cell viability data set, the distribution was found to be non-parametric. Therefore, a Kruskal-Wallis test followed by Dunn’s multiple comparisons test was performed. For the vessel deformation analysis, the association between proportional diameter change and vessel size was assessed by simple linear regression as described above. Differences with p < 0.05 were considered statistically significant.

## ACKNOWLEDGMENTS

The authors acknowledge support from the Swiss National Science Foundation (P2EZP2_199914 and P500PB_222131 to E.C.), the Ludwig Center at the MIT Koch Institute for Integrative Cancer Research (to E.C.), MathWorks (to M.A.F.), and the NIH National Cancer Institute (U54-CA261694 to R.D.K.). The authors thank the Koch Institute’s Robert A. Swanson (1969) Biotechnology Center for technical support, specifically Charlie Demurjian at the MIT BioMicro Center. The authors are also grateful to Kyle McKee for fruitful discussions about the mechanics of our microfluidic system. Figures were created in https://BioRender.com.

## AUTHOR CONTRIBUTIONS

M.A.F. and R.M.S. contributed equally to this work. Conceptualization, E.C.; Data curation, E.C.; Formal analysis, I.F. and E.C.; Funding acquisition, R.D.K. and E.C.; Investigation, M.A.F., R.M.S., R.N.C. and E.C.; Methodology, M.A.F., R.M.S., I.F., R.N.C. and E.C.; Project administration, E.C.; Resources, R.D.K.; Supervision, R.D.K. and E.C.; Visualization, M.A.F., I.F. and E.C.; Writing – original draft, M.A.F. and E.C.; Writing – review & editing, M.A.F., R.M.S., I.F., R.N.C., R.D.K. and E.C.

## DECLARATION OF INTERESTS

R.D.K. is a co-founder and a board member of AIM Biotech. He has also received research support from Amgen, AbbVie, Novartis, Daiichi-Sankyo, and Takeda. I.F. is an employee of Zimmer Biomet. These company relationships are unrelated to the content of this article. M.A.F., R.M.S., R.N.C., R.D.K., and E.C. are named inventors on a provisional patent application related to this work that was filed by MIT. The other authors declare no competing interests.

