## Supplementary material for "A mechanically actuated lung microvascular model reveals that breathing-like deformation enhances tumor cell extravasation": 2026_09_14_actuated_lung_vasculature_SI.pdf

[3] Independent researcher

[4] These authors contributed equally.

[5] Lead contact

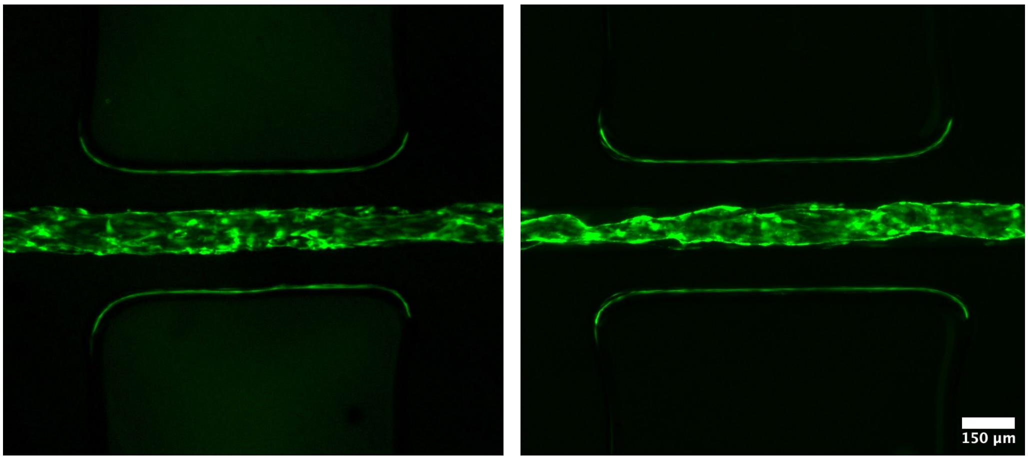


**Figure S1. Epifluorescence microscopy images of single large vessels formed by hPAECs in the 200-µm-wide channel.** The 200 µm width was insufficient to support the formation of microvascular networks. Instead, hPAECs (green) self-assembled into a single large vessel with little surrounding extracellular matrix available for tumor cell extravasation. Scale bar is 150 µm.
